# Human CD127 negative ILC2s show immunological memory

**DOI:** 10.1101/2023.10.29.564625

**Authors:** Laura Mathä, Lisette Krabbendam, Sergio Martinez-Høyer, Balthasar Heesters, Kornel Golebski, Chantal M.A Kradolfer, Maryam Ghaedi, Junjie Ma, Ralph Stadhouders, Claus Bachert, Lars O. Cardell, Zhang Nan, Gabriele Holtappels, Leanne C. Helgers, Theo B.H Geijtenbeek, Jonathan M. Coquet, Fumio Takei, Hergen Spits, Itziar Martinez-Gonzalez

## Abstract

Group 2 Innate Lymphoid Cells (ILC2s) serve as key players in type 2 immunity and contribute significantly to maintaining homeostasis and responding to inflammation. Notably, ILC2s are closely implicated in the development of allergic disorders like asthma. While previous research has demonstrated immunological memory in mouse ILC2s, it has remained unclear whether human ILC2s can acquire this form of memory. In this study, we demonstrate the persistence of CD45RO, a marker previously linked to inflammatory ILC2s, in resting ILC2s that have undergone prior activation. These cells concurrently reduce the expression of the canonical ILC marker CD127. Through in vitro experiments involving the isolation and stimulation of CD127-CD45RO+ ILC2s, we observed an augmented ability to produce cytokines and undergo proliferation. CD127-CD45RO+ ILC2s are found in both healthy and inflamed tissues and display a gene signature of cell activation. Memory ILC2s may play a significant role in chronic type 2 diseases, such as allergic asthma and atopic dermatitis.

**HIGHLIGHTS:**

– Inflamed and healthy tissues contain CD127-ILC2s with a gene signature of cell activation
– CD127-CD45RO+ ILC2s from healthy tissues show immunological memory features
– Mouse memory ILC2s downregulate CD127
– In vitro stimulation of naïve ILC2s generates memory CD127-CD45RO+ ILC2s

## INTRODUCTION

Group 2 Innate Lymphoid Cells (ILC2s) are abundant at barrier tissues in both mice and humans and are initiators of immune responses against allergens and parasites^1–3^. ILC2s are devoid of markers that define lineages such as T, B and myeloid cells but express IL-7Rα (CD127), IL-2Rα (CD25), the receptor for prostaglandin D2 (CRTH2)^4^ and inducible T cell costimulatory (ICOS) as well as the transcription factors GATA3^5,6^ and retinoic acid receptor-related orphan receptor alpha (RORA)^7^. ILC2s are mainly tissue resident^8^ and are activated by epithelial- or stromal-derived alarmins including the cytokines IL-33, IL-25 and thymic stromal lymphopoietin (TSLP)^9^. In response, they are capable of producing type 2 cytokines, including IL-4, IL-5 and IL-13, initiating an inflammatory cascade that leads to type 2 inflammation characterized by eosinophilia, mucus production and epithelial barrier leakage^10,11^. Consequently, ILC2s have been implicated in various allergic diseases and compared to healthy tissues higher numbers are found in inflamed tissues, such as skin of atopic dermatitis patients^12^ patients or nasal polyps from chronic rhinosinusitis with nasal polyps (CRSwNP) patients, compared to healthy tissues^13^.

We previously described immunological memory of ILC2s in a mouse model of lung inflammation^14^. Upon activation by IL-33 or allergens, ILC2s undergo an expansion phase marked by proliferation and IL-5 and IL-13 production. This is followed by a contraction phase where they stop producing cytokines but live for a long time. Upon secondary challenge by an unrelated allergen, previously activated ILC2s respond faster and more potently compared to naïve ILC2s, leading to enhanced inflammation. Whether human ILC2s can acquire memory functions remains unknown. We have recently shown that expression of CD45RO marks inflammatory ILC2s in humans^15^. Inflammatory ILC2s were highly activated and abundant in inflamed tissues such as nasal polyps from CRSwNP patients. CD45 is a receptor tyrosine phosphatase found on the surface of most blood cell types, including ILCs and T cells^16^. Whereas naive T cells express the CD45RA splicing isoform, activated and memory T cells exclusively express CD45RO. Inhibition of the activity of CD45 sensitizes ILC2s to cytokine production suggesting that this phosphatase is implicated in the regulation of activation of CD45RA+ ILC2s^15^.

We hypothesized that previously activated ILC2s could retain CD45RO expression and acquire memory features. Considering that humans are constantly exposed to extrinsic insults/stimuli, we suspected that memory ILC2s would be present in healthy individuals. CD127 is currently the canonical marker to identify ILCs in humans serving as a universal reference in the field for studying ILCs in various tissues and diverse disease contexts. Here, we show that human tissues contain a subset of CRTH2+ ILC2s that do not express CD127 and express high levels of CD45RO. CD127-CD45RO+ ILC2s are more responsive to epithelial alarmins than CD127+ ILC2s. Our data suggest that naïve ILC2s, upon activation, down-regulate CD127, switch from CD45RA+ to CD45RO+ and persist as memory ILC2s that are highly responsive to secondary challenges.

## Results

### CD127-ILC2s in inflamed tissues present an enhanced cell activation profile

Ma et al. recently published single cell RNA sequencing data on nasal polyps from asthma patients with CRSwNP^17^. ILC2s were isolated as live CD45+ CD3-Lin-CRTH2+ cells and clustered together (cluster 9, Supplementary Figure 1a) with a gene expression signature of *PTGDR2*, *GATA3*, *CD200R1*, *XCL1*, *IL9R* and *HPGDS* (Supplementary Figure 1b). Unsupervised reclustering with a higher resolution in this dataset further divided ILC2s into two clusters (Figure 1a and Supplementary Figure 1a), which were defined by differential expression of *IL7R* among other marker genes (p adj=0.0065) (Figure 1b and supplementary table 1). Therefore, we referred to these clusters as *IL7R* hi and *IL7R* low ILC2s. Both clusters showed equal expression of common ILC2 genes such as *PTGDR2, CD200R1* and *KLRB1* (Figure 1c); however, the *IL7R* low ILC2 cluster was enriched for the expression of type 2 cytokines such as *IL4*, *IL5* and *IL13*, alarmins receptors such as *IL1RL1* (IL-33 receptor) and *IL17RB* (IL-25 receptor) as well as *PTGS2, DUSP4, BATF, GATA3*, which suggested that this cluster holds an enhanced cell activation gene signature compared to *IL7R* hi ILC2s (Figure 1a, d).

**Figure 1.**
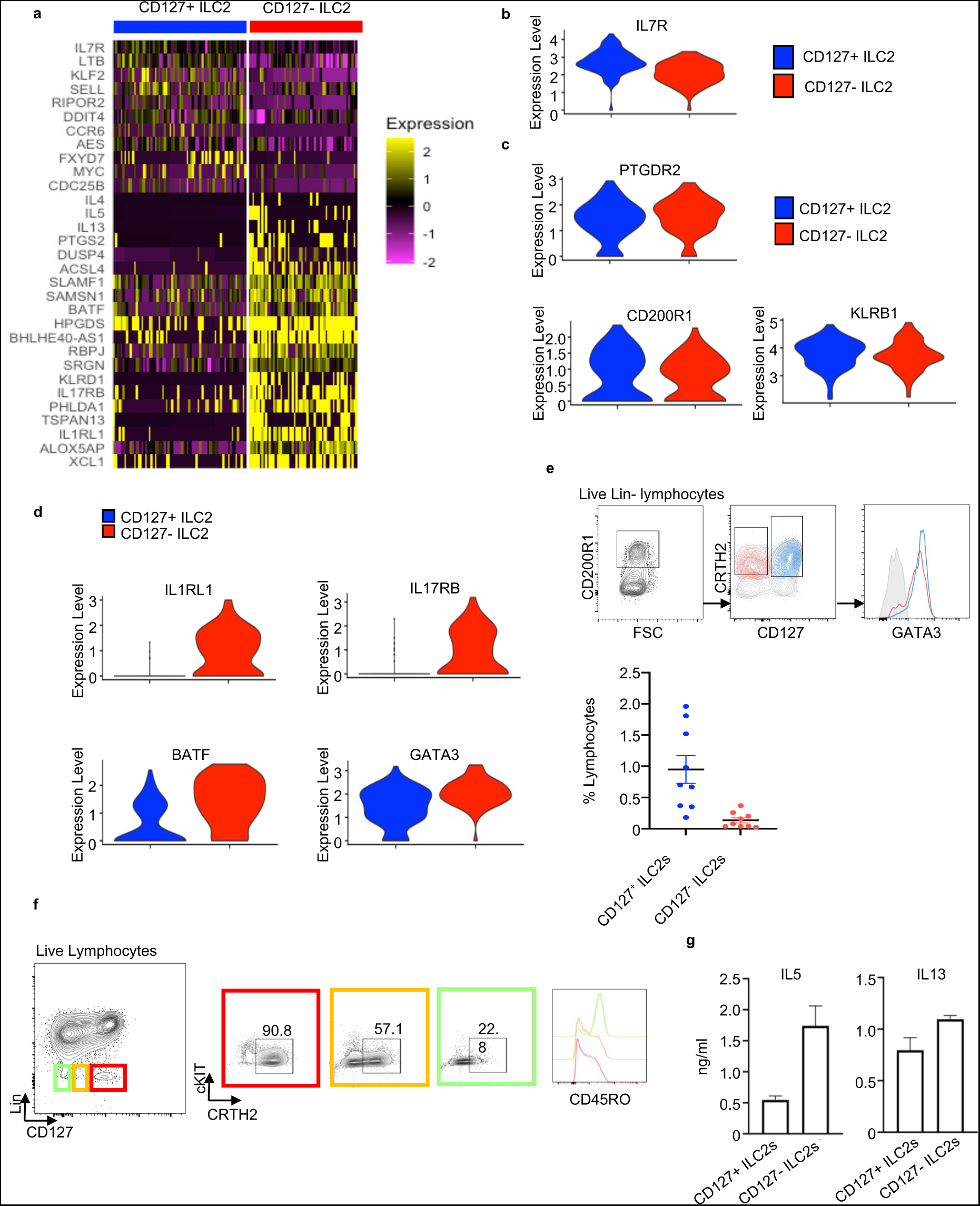
CD127-ILC2s in inflamed tissues present an enhanced cell activation profile. a) Heatmap of top marker genes from the CD127+ and CD127- ILC2 clusters observed in the scRNAseq dataset from nasal polyps from CRSwNP patients^17^. b) Violin plot of *IL7R* expression in the CD127+ and CD127- ILC2 clusters. C-d) Violin plots of the indicated genes CD127+ and CD127- ILC2 clusters. e) Quantification and GATA3 expression of CD127+ and CD127- ILC2s in fresh nasal polyps from CRSwNP patients analyzed by flow cytometry. f) CD45RO expression on CD127+ and CD127- ILC2s f in fresh nasal polyps from CRSwNP patients analyzed by flow cytometry. g) ELISA quantification of IL-5 and IL-13 levels in the supernatant of in vitro stimulated CD127+ and CD127- ILC2s isolated from nasal polyps with IL2+IL7+IL33+TSLP for 7 days. ScRNAseq data analysis was performed using Seurat.

We also reanalyzed published scRNA sequencing data from punch biopsies patients with AD^18^ (Supplementary Figure 1c). This analysis revealed that the ILC2 clusters (0-3) can be further subdivided into two groups: one primarily characterized by the expression of genes typically associated with naïve ILC2s, including *IL7R*, and the other marked by the expression of genes indicative of activated ILC2s, such as *IL5, IL13, IL1RL1, and IL17RB,* but with lower levels of *IL7R*. Consequently, *IL7R*-low ILC2s, which are enriched in cytokine and effector mRNA expression, were notably present in nasal polyp tissue and AD lesions (Supplementary Figure 1d).

To confirm the presence of *IL7R* low ILC2s in vivo, we performed further analysis of nasal polyp samples by flow cytometry. In our analyses, we identified a subset of CD127-ILC2s, albeit present at a lower frequency than CD127+ ILC2s (Figure 1e). Furthermore, the expression of the surface marker CD45RO was higher in CD127-ILC2s compared to CD127+ ILC2s (Figure 1f).

Given the negative correlation observed between *IL7R* expression and a gene signature of cell activation in ILC2s, we tested the responsiveness of CD127-ILC2s. Purified CD127-ILC2s from nasal polyps showed an enhanced response to cytokine stimulation by producing higher amounts of type 2 cytokines compared to CD127+ ILC2s (Figure 1g).

Altogether, these data suggest that tissues with a type 2 inflammation signature, including nasal polyps and AD samples, contain CD127-ILC2s characterized by an enhanced cell activation gene signature and greater ability to respond to stimuli.

### CD127-ILC2s are present in healthy tissues

The observation of two distinct ILC2 cellular states in inflamed tissues led us to ask whether healthy tissues also contained CD127-ILC2s. To this end, we purified CD45+ Lin-cells from healthy human dermis directly after surgical removal and analyzed them by single cell RNA sequencing (scRNAseq). We performed plate-based scRNAseq^19^ to be able to back trace protein expression by using index sorting at the time of the purification. The identification of a CD127-ILC2 population called for another surface marker to identify ILC2s within Lin-cells. CD200R1 has been shown to be expressed on all human and mouse ILC subsets in PB, intestine and tonsil but showed the highest expression on ILC2s^20^. We observed that dermal CD127+ ILC2s also highly expressed CD200R1, in contrast to ILC1 and ILC3s (Supplementary Fig 2a), and CD200R1 expression on ILC2s did not change upon in vitro stimulation (Supplementary Fig 2b). To enrich for ILCs within the Lin-compartment and optimize our scRNAseq approach, equal numbers of cells were sorted in 4 distinct groups: CD127+ CD200R1+, CD127+ CD200R1-, CD127-CD200R1+ and CD127-CD200R1-(Supplementary Figure 2c). CD45+ CD3+ CD45RO+ T cells were added as a reference. From 1583 cells sequenced, high-quality transcriptomes were obtained from 1449 cells and further analyzed. Dimension reduction using UMAP revealed 10 clusters (Supplementary Figure 2d), including ILCs and some unexpected populations such as endothelial cells, fibroblasts, and dermal dendritic cells. ILCs, which expressed *KLRB1*, *IL7R*, *AREG*, *TNFRSF18* and *CD69* and were negative for *CD3D,* were divided into 3 clusters: ILCa, ILCb and ILCc (Supplementary Figure 2e). Analysis of the distribution of *IL7R* expression within the ILC clusters showed that cluster ILCa expressed lower levels of *IL7R* mRNA compared to ILCb and ILCc (Figure 2 a-b). This was corroborated by the level of CD127 protein expressed in these ILC populations (Figure 2c).

**Figure 2.**
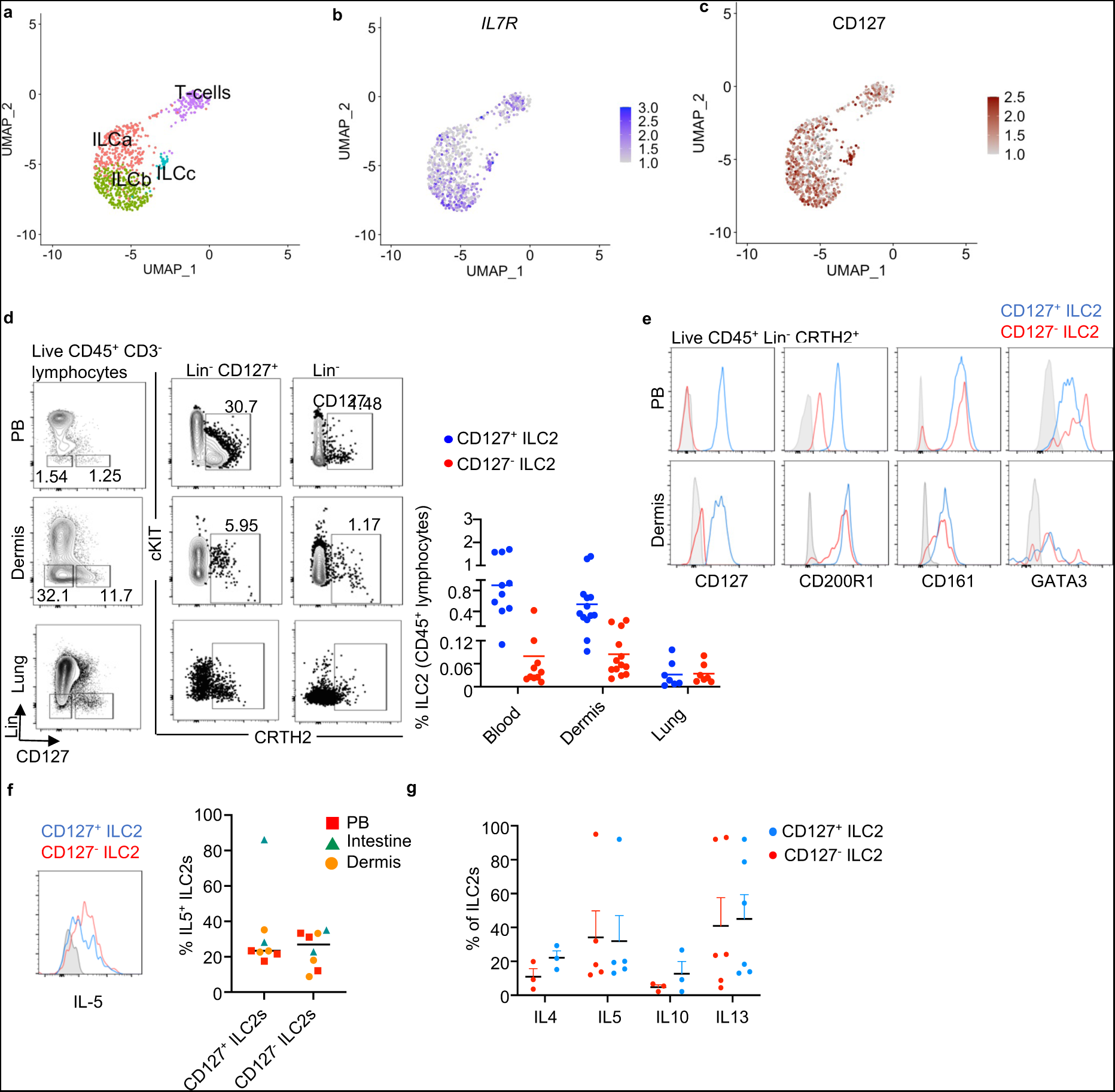
CD127- ILC2s are present in healthy tissues. Single cell RNA sequencing was performed in healthy human dermal cells (see gating strategy on Supplementary Figure 2c). a) UMAP visual representation of the ILC and T cell clusters obtained after filtering out non relevant cell-types in the Lineage negative compartment. b) Expression level of *IL7R* plotted in blue on the ILC and T cell clusters. The intensity of color indicates the expression level. c) Index sorting overlay of CD127 expressing cells on the UMAP clustering. Brown dots indicate CD127 expressing cells. The intensity of color indicates the expression level. d) Flow cytometry analysis of CD127+ and CD127- ILC2s in human fresh PB, dermis and lung. Lin: CD1a, CD3, CD5, CD8, CD11c, CD14, CD16, CD19, CD34, CD94, CD123, FceR1a, TCRαβ, TCRgd, BDCA2. e) Flow cytometry measurement of the expression of CD127, CD200R, CD161 and GATA3 on CD127+ and CD127- ILC2s from PB and dermis. f) CD127+ and CD127- ILC2s were isolated from fresh PB, dermis and adult intestine and in vitro stimulated with IL-2 + IL-7 + IL-33 (IL-25 for dermis) + TSLP for 5 days. Intracellular staining of IL-5 was performed and analyzed by flow cytometry. g) Quantification of the expression of the indicated cytokines in the CD127+ and CD127- ILC2s isolated from PB and stimulated with IL-2 + IL-7 + IL-33 + TSLP for 5 days. Data from e is representative of at least three donors from more than three independent experiments. Symbols represent individual donors; bars indicate mean values. ScRNAseq data analysis was performed using Seurat.

Flow cytometry analysis of several tissues including PB and dermis confirmed the presence of CD127-ILC2s albeit in lower frequency than CD127+ ILC2s (Figure 2d). These CD127-ILC2s presented a typical ILC2 phenotype, expressing CD200R1, CD161 and GATA3 (Figure 2e). Remarkably, we also found ILC2s as CD127-ILC2s in the adult intestine, typically considered rare in humans despite their abundance in the fetal intestine (Supplementary Figure 2f-g).

Imaging flow cytometry (ImageStream) analysis^21^ confirmed the lymphocytic morphology of CD127-ILC2s and showed low amounts of CD127 on the cell surface (Supplementary Figure 2h). Quantitative PCR of *IL7R* showed that ILC2 from several tissues expressed lower levels of *IL7R* in CD127-cells compared to CD127+ ILC2s, in agreement with the dermis scRNA sequencing data (Supplementary Figure 2i), correlating mRNA and protein expression and therefore ruling out the possibility of technical artefacts introduced by tissue processing or immunophenotyping during our analyses. These data suggest that CD127-ILC2 do not constitute a separate lineage independent of IL-7 signaling. Although these cells express some *IL-7R* transcripts, they appear negative in flow cytometry analyses and therefore we will be referring to them as CD127-ILC2s.

To determine the functionality of CD127-ILC2s, we purified Lin-CD127-CRTH2+ cells and Lin-CD127+ CRTH2+ ILC2s from various tissues and stimulated them *in vitro* with an ILC2 activating cytokine cocktail. Upon 5 days of stimulation, Lin-CD127-CRTH2+ cells became positive for intracellular IL-5 staining (Figure 2f). Furthermore, other ILC2 related cytokines including IL-4, IL-10 and IL-13 were detected in PB CD127-ILC2s (Figure 2g).

To validate the identity of CD127-ILC2s, we conducted single-cell cloning assays. We cultured Lin-CD127+ and CD127- from individual cells, supported by mouse stromal cells (OP9-DL1), and in the presence of a cytokine cocktail consisting of IL-2, IL-7, IL1β, and IL-23, known to promote the expansion of all ILC subtypes^20,22^. After two weeks, we analyzed the expanded clones, with 23 originating from the CD127+ ILCs and 21 from the CD127-ILCs, for cytokine expression using flow cytometry (Supplementary Figure 2j). 26% of the CD127+ and 23.8% of the CD127-ILC clones were true ILC2s (producing IL-5, IL-4, IL-13 and/or IL-17F) (Supplementary Figure 2k-l). Additionally, we identified ILC1 and ILC3 clones, as well as other clones with a plastic ILC phenotype (Supplementary Figure 2m). Consequently, it is evident that the Lin-CD127- compartment indeed contains functional and bona fide ILC2s. Altogether, our data show that some functional ILC2s present in tissues and in circulation of healthy individuals do not express CD127 on their cell surface.

### CD45RO expression is enriched within the CD127-ILC2s in healthy tissues

To determine differentially expressed genes between the ILC clusters in the dermis, we performed a non-parametric Wilcoxon rank sum test. In addition to the significantly lower expression of *IL7R* in ILCa compared to ILCb and ILCc, this cluster was characterized by the expression of activation related genes including *CD69*, *DUSP1*, *FOS*, *JUN*, *KLF6*, *NFKBIA* and *NR4A1* (Figure 3a). Accordingly, pathway enrichment analysis showed a gene signature for MAPK activation as the top pathway represented in the ILCa cluster (Figure 3b). Index sorting data showed enrichment of CD45RO+ cells within the ILCa cluster (Figure 3c), consistent with our previous analyses in nasal polyps. Importantly, flow cytometry analysis confirmed that CD127-ILC2s expressed CD45RO and were negative for CD45RA, in contrast to CD127+ ILC2s (Figure 3d). Therefore, CD127-CD45RO+ ILC2s are not only present in inflamed tissues but also in healthy tissues, carrying a gene signature of cell activation.

**Figure 3.**
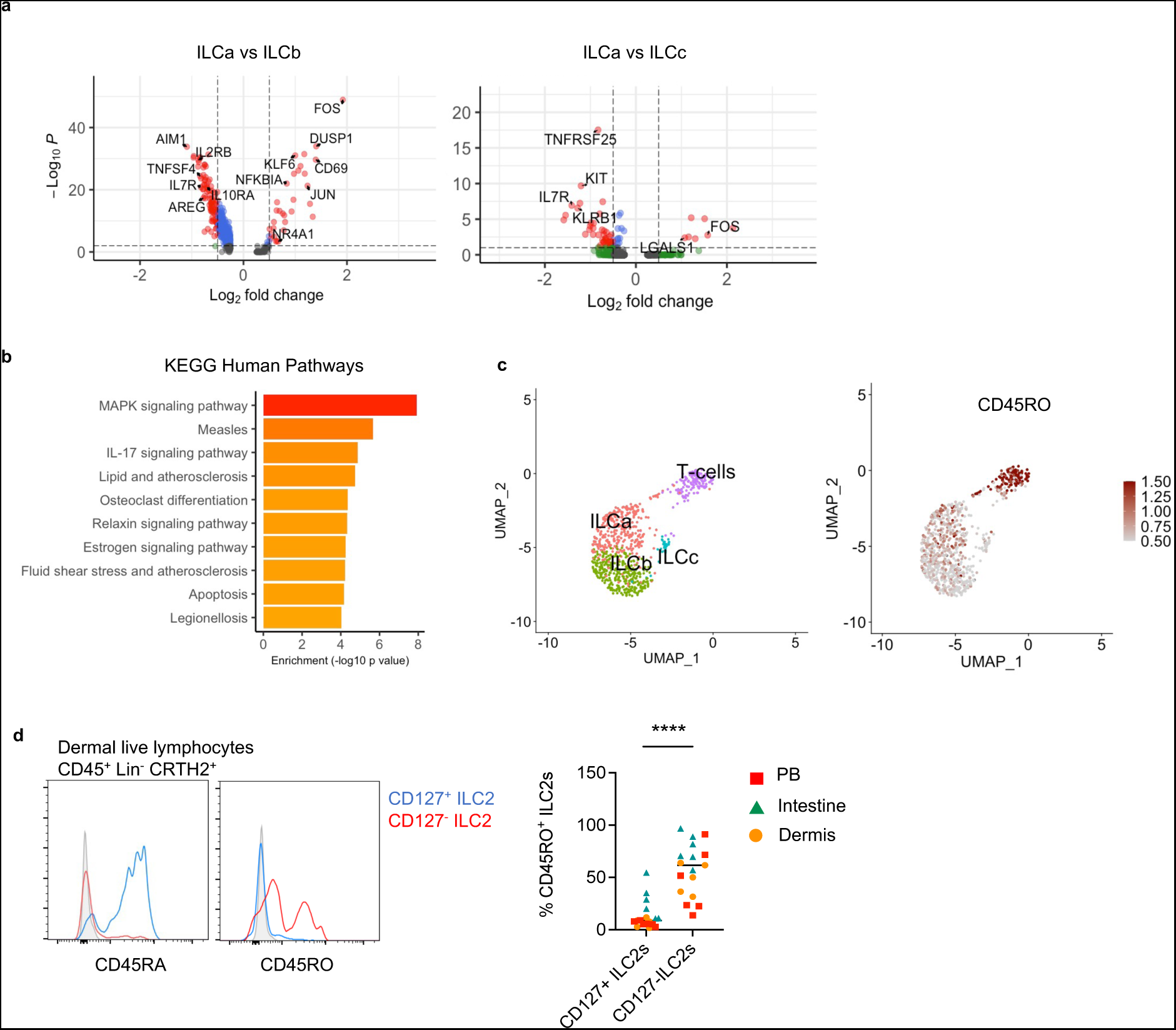
CD127- ILC2s express high levels of CD45RO in healthy tissues. a) Volcano plots showing differential gene expression between the ILC clusters. Statistic significant genes (marked in red) were calculated using a non-parametric Wilcoxon rank sum test (p<0.001, log fold change <0.5). Red color marks genes above the p value and fold change thresholds. b) Pathway enrichment based on differential gene expression for cluster ILCa compared to ILCb. c) Index sorting overlay of CD45RO expressing cells on the UMAP re-clustering. Red dots indicate CD45RO expressing cells. d) Flow cytometry measurement of CD45RA and CD45RO expression on CD127+ and CD127- ILC2s from dermis and quantification of the percentage of CD45RO expression on CD127+ and CD127- ILC2s from the indicated tissues. Symbols represent individual donors; bars indicate mean values. ****P < 0.0001 (Paired t test). ScRNAseq data analysis was performed using Seurat.

### CD127-CD45RO+ ILC2s from healthy donors are more responsive than CD127+ CD45RA+ ILC2s

We have previously observed that mouse memory ILC2s are characterized by increased transcription of genes related to ILC2 activation^14^. Therefore, we hypothesized that human CD127-CD45RO+ ILC2s in healthy tissues, marked by an activation gene signature, could have been previously activated ILC2s that remained in the tissue. In fact, it is conceivable that healthy tissues, especially surface barriers, contain previously activated or experienced ILC2s as our body is constantly exposed to extrinsic stimuli that can activate ILC2s. In agreement, previously activated tissue resident memory T cells are marked by the expression of CD45RO^16^. To investigate the potential immunological memory properties of CD127-CD45RO+ ILC2s, we purified CD127-CD45RO+ and CD127+ CD45RA+ ILC2s from healthy PB and dermis by FACS (Figure 4a). To study the rapid response typically found in immune memory cell, we performed a 24h *in vitro* stimulation with an ILC2 stimulatory cytokine cocktail (IL-2 + IL-7 + IL-33 + TSLP and IL-2 + IL-7 + IL-25 + TSLP for PB and dermis, respectively). CD45RO+ ILC2s produced significantly larger amounts of IL-5 than CD45RA+ ILC2s (Figure 4b and Supplementary Figure 3a). Of note, isolated CD45RO+ ILC2s did not produce cytokines when cultured with IL-2+IL-7 (Supplementary Figure 3b), suggesting that these cells were not actively producing cytokines in the tissue. Finally, CD45RO+ ILC2s from dermis showed increased proliferation in response to a 48h stimulation with IL-2 + IL-7 + IL-25 + TSLP compared to CD45RA+ ILC2s (Figure 4c). As mentioned earlier, there might be traces of CD127 on the cell surface of CD127-ILC2s, enabling TSLP to activate them.

**Figure 4.**
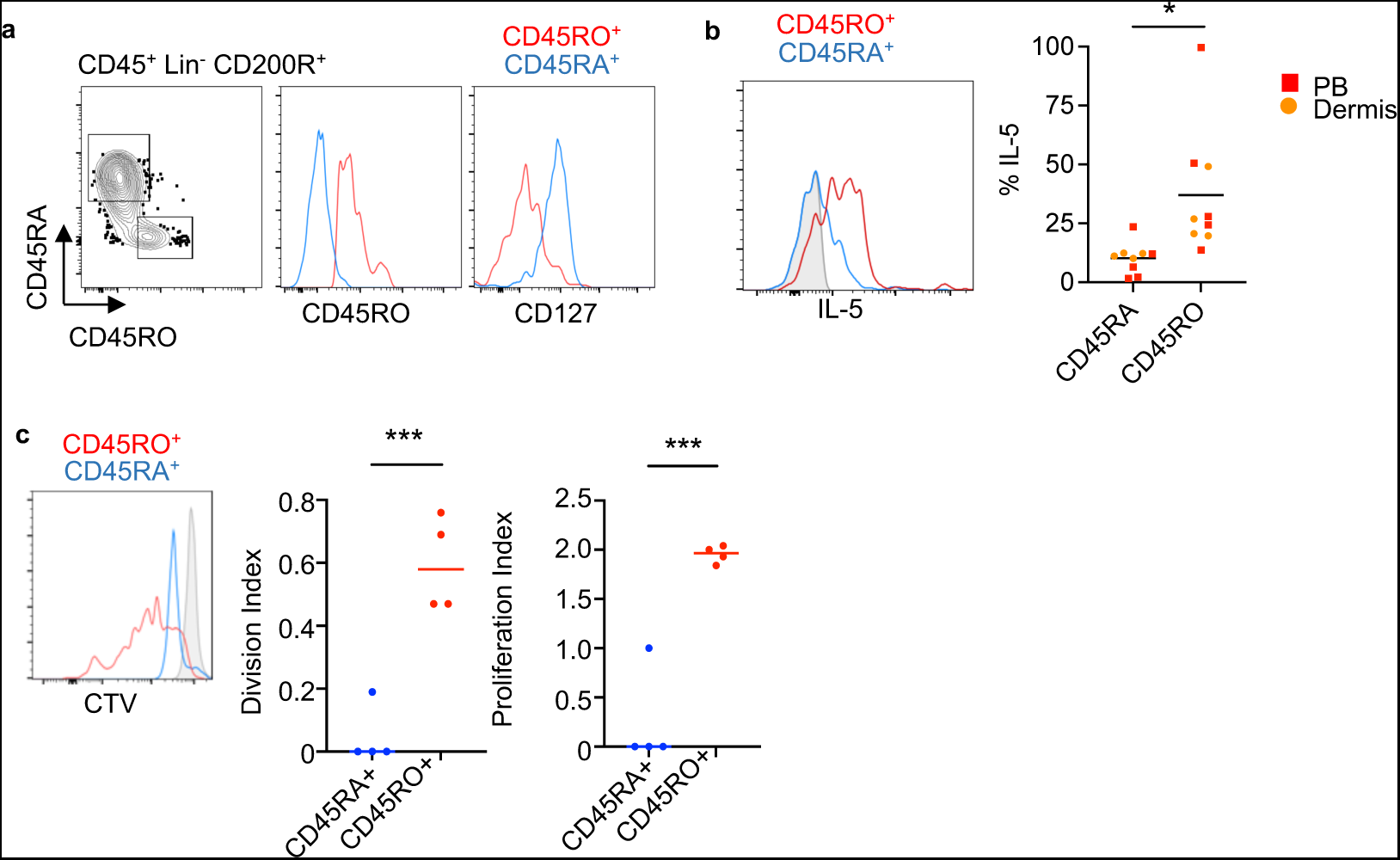
CD45RO+ ILC2s from healthy donors are more responsive than CD45RA+ ILC2s. a) Sorting strategy of CD45RA+ (CD127+) and CD45RO+ (CD127-) ILC2s isolated from healthy human dermis. An equal number of cells were plated in a round bottom 96 well plate. b) Histogram and quantification showing IL-5 expression on stimulated CD45RA+ and CD45RO+ ILC2s from the indicated tissues. PB ILC2s were stimulated with IL-2 + IL-7 + IL-33 + TSLP and dermal ILC2s with IL-2 + IL-7 + IL-25 + TSLP for 24 hours. c) CTV staining of in vitro stimulated CD45RA+ and CD45RO+ ILC2s purified from healthy dermis as in a), with the same cytokine cocktail as in b) for 48 h. Symbols represent individual donors; bars indicate mean values. *P < 0.05, and ***P < 0.001, (Paired t test).

In summary, these findings indicate that CD127-CD45RO+ ILC2s are present in both the bloodstream and as resident cells within tissues. Moreover, they exhibit a more rapid and robust cytokine production capacity when compared to CD127+ CD45RA+ ILC2s, strongly implying their identity as memory ILC2s.

### CD127-ILC2s are present in mouse tissues

To investigate whether CD127-ILC2s are present in mouse tissues, we analyzed naïve B6 lungs, small intestine, skin, and liver by flow cytometry and identified CD127+ and CD127- populations within respective ILC2 gates (Figure 5a). In the lung, CD127+ and CD127-ILC2 populations expressed similar levels of ILC2 related surface markers and transcription factors including ST2, CD25, KLRG1 and GATA3, but not Tbet and RORγt, demonstrating phenotypic similarities between CD127-ILC2s and CD127+ ILC2s (Figure 5b). To further confirm an ILC2 lineage identity, we analyzed the RORα-lineage tracer Rora-YFP mice, in which almost 90% of ILC2s express YFP^23^. CD127+ and CD127-subsets of ILC2s defined by Lin-RORγt-Tbet-Thy1+ GATA3+ cells expressed YFP and ILC2 related surface markers, confirming that they belong to the RORα+ ILC2 lineage (Figure 5c). Intracellular staining of the lungs after IL-33 treatment demonstrated that CD127+ and CD127-ILC2s have a similar ability to produce IL-5 and IL-13, indicating that the CD127-cells are functional ILC2s (Figure 5d). Altogether, these results demonstrate that CD127-ILC2s in mice phenotypically and functionally resemble conventional CD127+ ILC2s, suggesting that they belong to the ILC2 lineage. Furthermore, their existence in mice supports the identification of the previously unappreciated CD127- population in humans.

**Figure 5.**
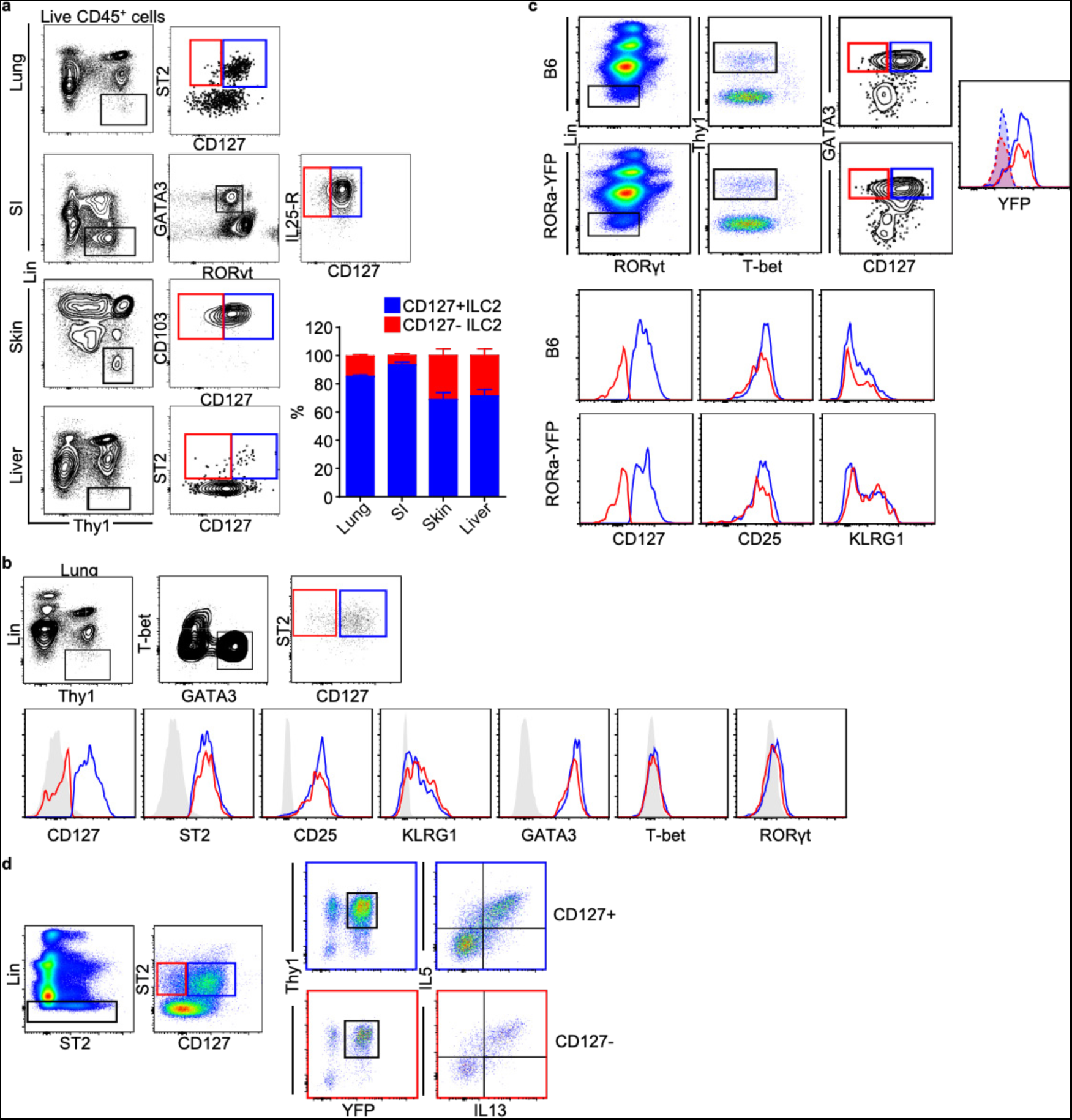
CD127- ILC2s are present in mouse tissues. a) Gating strategies used to quantify CD127^+^ (blue) and CD127^-^ (red) ILC2s in various tissues (plots on the left) and quantification of the two subsets (graph on the right). b) Phenotypic characterization of CD127^+^ (blue) and CD127^-^ (red) ILC2 in mouse lungs. c)Identification and phenotypic characterization of CD127^+^ (blue) and CD127^-^ (red) ILC2s in B6 and Rora-YFP mouse lungs. d)IL-5 and IL-13 expression in CD127^+^ and CD127^-^ ILC2 subsets from Rora-YFP mouse lungs after IL-33 stimulation. Data represented are mean ± SEM. n=4; 1 experiment (skin), n=6; 2 independent experiments (small intestine), n=13; 4 independent experiments (lung and liver).

### Memory ILC2s in mice present reduced expression of CD127

We have previously shown the role of memory ILC2s in a mouse model of allergic lung inflammation^14^. To investigate whether memory ILC2s in mice also modulated the expression of CD127, we analyzed ILC2s in the lungs of naïve, effector (72h after IL-33 intranasal administration) and memory (1 month after IL-33 intranasal administration) mice (Figure 6a). The percentage of CD127-ILC2s was similar in naïve and effector lungs, while it was significantly increased in memory mouse lungs (Figure 6b). Upon challenge with papain, most of the cytokines were derived from the CD127-ILC2s rather than CD127+ ILC2s in the memory mice (Figure 6c). These findings indicate that memory ILC2s in mice show an enrichment within the CD127- subset, mirroring our observations in human ILC2s.

**Figure 6.**
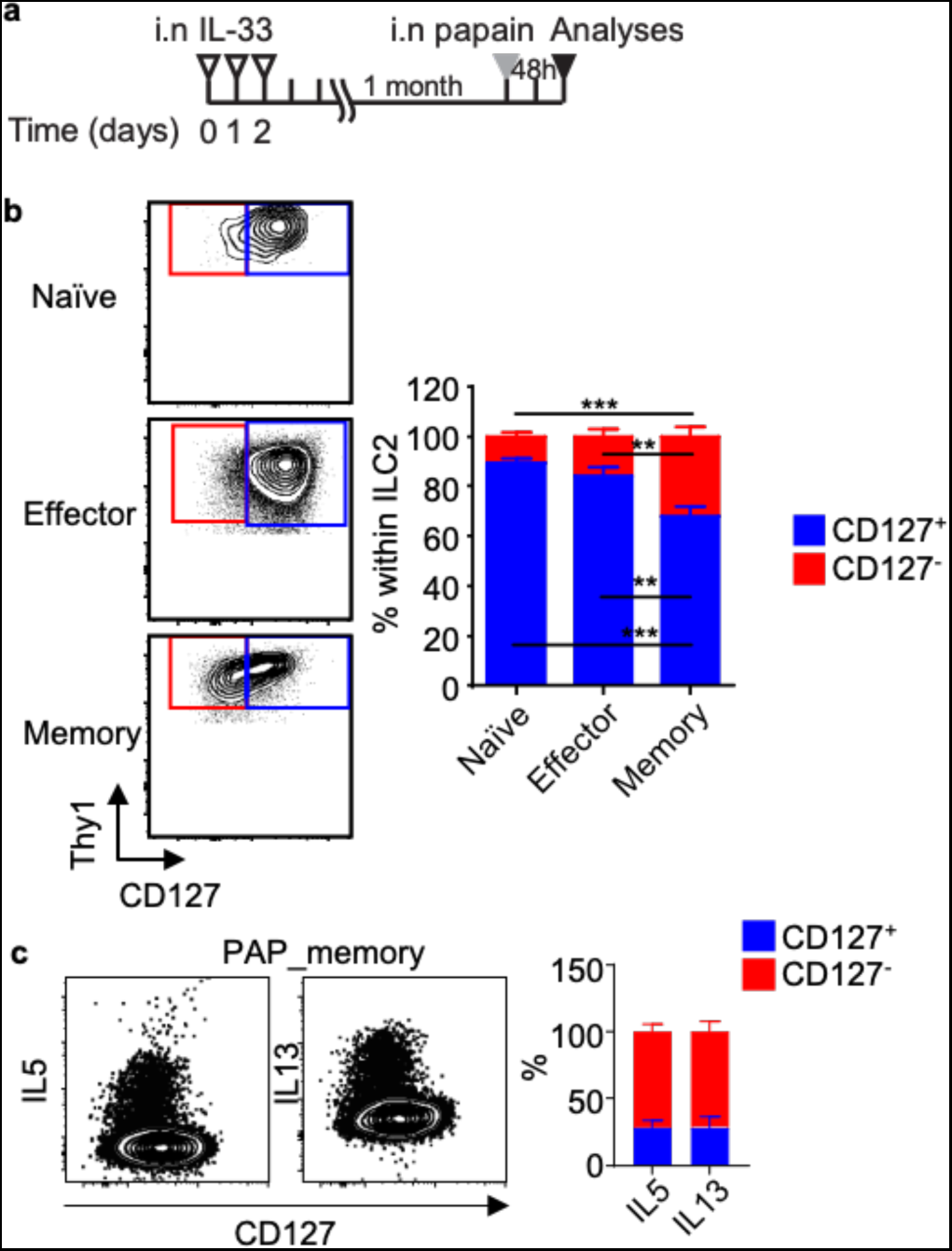
Memory ILC2s in mice present reduced expression of CD127. a) Mice were treated with 3 consecutive days of i.n. IL-33 injections and naïve, effector (72 hours after the last injections) and memory (1 month after the last injections) lungs were harvested. b) CD127+ (blue) and CD127- ILC2s (red) were analysed by flow cytometry in naïve, effector and memory cells. c) The dot plots show IL-5 and IL-13 producing ILC2s, while the graph shows the percentages of IL- 5^+^ and IL-13^+^ ILC2s in the CD127^+^ (blue) and CD127^-^ (red) compartments. Data presented are mean ± SEM. n=6; 2 independent experiments. A paired two-tailed t test was used to determine statistical significance, with a P value <0.05 being significant. *P<0.05, **P<0.01, ***P<0.001.

### In vitro stimulation of CD127+ ILC2s generated highly responsive CD127-CD45RO+ ILC2s

We have observed that stimulation of human ILC2s induces downregulation of CD127 expression. To investigate whether CD127+ ILC2s could give rise to memory CD127-CD45RO+ ILC2s, we sorted CD127+ CD45RA+ ILC2s from several tissues including dermis, peripheral blood (PB), intestine, tonsil and cord blood (see a complete gating strategy from PB as an example in Supplementary Figure 4a) and in vitro stimulated them with IL-2 + IL-7 + IL-33 + TSLP for 7 days. The combination of IL-2 and IL-7 was used to maintain ILC2s alive as control (Figure 7a). After one week of stimulation, the cells were washed and fresh media containing only IL-2 + IL-7 were added to induce a resting state. Under these conditions, we observed that human ILC2s expanded in numbers upon activation, followed by a contraction phase upon alarmin withdrawal, during which a small ILC2 population survived and remained alive for up to 1 month (Figure 7b). Primed ILC2s showed increased survival than control ILC2s at day 21 (71.5±3.6 vs 38.9±5.5 % of viable cells, respectively) (Figure 7c). Following stimulation, ILC2s underwent a phenotypic change characterized by the increased expression of CD45RO and the reduced expression of CD127 (refer to Supplementary Figure 4b). Throughout the expansion phase, ILC2s continued to release IL-5 and IL-13 (see Figure 7d-e). In line with previous observations, cytokines were mostly produced by CD127-CD45RO+ ILC2s (Figure 7f). ILC2s retained the expression of CD45RO during the culture’s resting phase even when they were no longer actively producing cytokines (Figure 7g). To determine whether these previously activated ILC2s were more responsive to a secondary challenge compared to unprimed cells, we added IL-33 to the culture for a brief period (16 h) on day 30 and analyzed cytokine production by flow cytometry. Primed ILC2s strongly responded to the IL-33 challenge by producing IL-5 and IL-13, in contrast to a negligible response observed in unprimed ILC2s (maintained with only IL-2 + IL-7 for 30 days) (Figure 7h and Supplementary Figure 4c).

**Figure 7.**
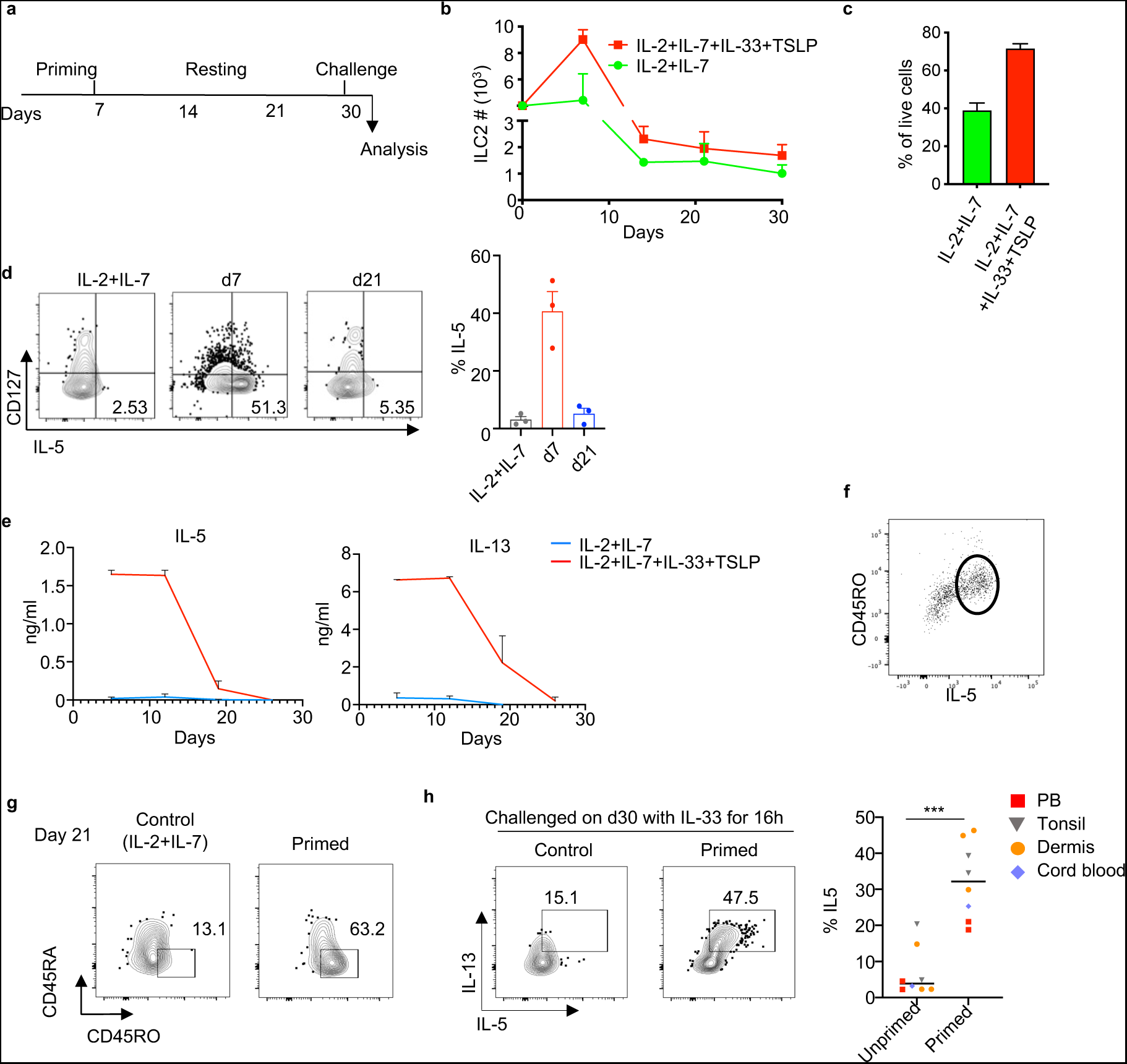
In vitro generated human CD45RO+ ILC2s showed immunological memory properties. a) Scheme of in vitro priming and challenge of purified CD127+ CD45RA+ ILC2s isolated from healthy PB or dermis. ILC2s were either primed with IL-2+IL-7+IL-33+TSLP or maintained in IL-2+IL-7 as control for 7 days. Then, wells were washed, and media was refreshed with IL-2+IL-7 every week. ILC2s were challenged with IL33 or PBS as control on day 30 and were stained and analyzed by flow cytometry on day 31 after PMA+Ionomicyn+Golgistop incubation for 3h. b) Time course analysis of number of ILC2s during the in vitro culture experiment as in a. c) % viable cells in the cultured ILC2s at the different time points. d) Flow cytometry measurement of CD127 and IL-5 expression on cultured ILC2s at the indicated time points of the culture. e) Time course of IL-5 and IL-13 levels in the supernatant of the cultured ILC2s quantified by ELISA. f) Expression of CD45RO and IL-5 of primed ILC2s on day 7. g) Flow cytometry measurement of CD45RA and CD45RO of cultured ILC2s on day 21. h) Flow cytometry measurement of IL-5 expression on unprimed (IL-2+IL-7) and primed (IL-2+IL- 7+IL-33+TSLP) ILC2s receiving IL-33 challenge. Data from f) and g) are representative of at least three donors from more than three independent experiments. Symbols represent individual donors; bars indicate mean values. ***P < 0.001, (Paired t test).

Interestingly, in vitro generated CD127-CD45RO+ ILC2s regained CD127 expression when IL-7 was withdrawn from the culture (Supplementary Figure 4d-e), suggesting that CD127 expression can be regulated. Nevertheless, they still maintained the expression of CD45RO. This observation prompted us to examine whether CD127-ILC2s isolated from healthy tissues could regain the expression of CD127. We isolated dermal CD127-ILC2s (and CD127+ ILC2s as control) and cultured them for 24h in the presence of IL-2 (Supplementary Figure 4f). The majority of the sorted CD127-ILC2s upregulated the surface expression of CD127 (Supplementary Figure 4g). Of note, we used two different fluorescent labels for the CD127 antibody at the time of the sorting and after the overnight culture. In conclusion, human CD45RA+ ILC2s become CD127-CD45RO+ ILC2s upon stimulation. Once they return to a resting state, they retain CD45RO expression and are intrinsically more responsive to restimulation compared to unprimed ILC2s, suggesting they have acquired immunological memory. Unlike CD45RO, the expression of CD127 in memory ILC2s seems to be adaptable to external signals.

## DISCUSSION

Here, we characterize immunological memory within human ILC2s as their ability to “recall” prior activation events, manifesting as a heightened response to secondary stimuli compared to the initial ones. Memory ILC2s are defined as CD127-CD45RO+ cells. In comparison with CD45RA+ ILC2, they exhibit increased proliferation rates and produce larger quantities of type 2 cytokines when exposed to epithelial alarmins. Expression of CD45RO in ILC2s found in healthy tissues indicates that they have previously undergone activation events. It is worth noting that memory ILC2s may also exist within the CD127+ CD45RO+ ILC2 subset, as observed in nasal polyps from patients with CRSwNP suggesting that downregulation of CD127 is a dynamic process in the development of memory ILC2s. However, in healthy tissues like the skin, CD127+ ILC2s predominantly express CD45RA.

ILCs rely on IL-7 for their development^24^ and therefore express the receptor for this cytokine, CD127. In fact, CD127 has been used as a critical marker to identify human ILCs^25^. Whereas CD127+ ILC2s are readily detected in most human fetal and adult tissues, they are rare in adult intestine^26^. We have now found ILC2s within the CD127 negative compartment, suggesting that in the human adult intestine, ILC2s mostly lack CD127. It may be possible that CD127+ ILC2s, which are abundant in the fetal intestine, modify their phenotype by downregulating the expression of CD127 after birth following interactions with the gut microbiome or products from food digestion. Interestingly, CD127+ ILC2s are abundant in the intestine of mice. Whether this is a consequence of being housed in specific pathogen free (SPF) conditions should be addressed in future studies.

Memory ILC2s are characterized by a transcriptional reprogramming towards cell activation and enriched for MAPK pathway members, such as *FOS* and *JUN*, which are downstream effectors of p38 activation. Importantly, p38 MAPK has been shown to be a central pathway of memory Th2 cells to respond to IL-33 in mouse and human^27^. It is likely that activated ILC2s downregulate CD127 expression as they become CD45RO+ memory cells. In this context, it is tempting to speculate that memory ILC2s remain in tissues by relocating into IL-7 rich environments, therefore explaining the downregulation of CD127. We cannot exclude the possibility that some memory ILC2s have stably downregulated CD127, especially in the adult human intestine where CD127+ ILC2s are rarely present. Zwijnenburg et al.^28^ have recently suggested that T cell differentiation occurs as a continuum of cellular states rather than transitioning between discrete subsets. They identified CX3CR1 as a graded marker for functionally distinct states of human antigen experienced CD8+ T cells. It is likely that naïve, activated and memory ILC2s can also be identified as a continuum of cellular states rather than distinct subsets. The inverse correlation of CD45RO and CD127 expressions on ILC2s in the nasal polyps indicates that CD127 expression gradient on ILC2s may reflect the broad spectrum of cell state on ILC2s.

Downregulation of CD127 on IL-33 activated mouse ILC2s has been previously suggested^29,30^. We now show that this low expression of CD127 on ILC2s is maintained as they acquire immunological memory and that CD127- memory ILC2s are the major producers of IL-5 and IL-13 upon re-challenge. Naive mice also have CD127-ILC2s, suggesting that these cells have a history of being activated. Indeed, previous reports^31^ have shown that after weaning a transient wave of IL-33 in peripheral tissues prime ILC2s which then increase expression of *Nr4a1*, associated with a heightened responsiveness of ILC2s. Interestingly, our data shows that *NR4A1* expression is significantly higher in skin CD127-ILCs from healthy donors as compared to CD45RA+ILC2s.

Finally, we have also identified Lin-CD127-CRTH2-CD200R+ CD45RO+ cells within the cKIT+ and cKIT-gates (data not shown), suggesting the existence of memory ILC1 and ILC3. Of note, mouse memory ILC3 has been recently described^32^. It may be possible that the Lin-CD127-CRTH2-CD200R+ CD45RO+ population also contains memory ILC2s which have downregulated CRTH2 as it has been shown that CRTH2-ILCs from human blood and lung cells can produce IL-5^33^.

Our results highlight the need for a re-evaluation of the established gating strategies used to identify and isolate human ILCs in various tissues, as the ability of ILC2s to regulate the expression of CD127, a well-recognized marker for human ILCs, has become evident. Instead, a combination of CD127 with the expression of CD45RA and CD45RO may offer insights into the cellular state of ILC2s.

Alternatively, the CD200R1 marker, in conjunction with CD161, emerges as a promising candidate that could enhance our gating strategy for defining human ILC2s within tissues. Furthermore, the discovery of highly responsive CD127-CD45RO+ ILC2s in inflamed nasal polyps from patients with chronic rhinosinusitis with nasal polyps (CRSwNP) suggests the potential relevance of memory ILC2s in this and other type 2 diseases. In line with this, our recent observations indicate that the presence of CD45RO+ ILC2s correlates with a quicker clinical response among patients with chronic CRSwNP receiving treatment with the anti-IL-4Ra antibody dipulimab^34^. Immunological memory of ILC2s may have important implications in allergic diseases especially in the so called non-allergic asthma^35^, as in contrast to T cells, ILC2s respond in an antigen non-specific manner. Therefore, it may explain why some allergic patients react to unrelated allergens.

## ACKNOWLEDGEMENTS

The authors thank Thomas Van Wingen for experimental support. IMG was financially supported by Marie Sklodowska-Curie grant No 798927 and Karolinska Institute. The project was financially supported by the advanced ERC grant 341038 to HS, the type 2 innovation grant from Sanofi Genzyme, the Swedish Research Council and the Knut and Alice Wallenberg foundation and Karolinska Institute to IMG.

## AUTHOR CONTRIBUTIONS

IMG conceived the study, designed, and performed the experiments, interpreted the data, wrote the manuscript, supervised the study, and acquired funding. HS acquired funding, supervised part of the study in humans and edited the manuscript. LM, LK and. SMH performed experiments, interpreted the data, and co-wrote the manuscript. SMH, BH and JC performed bioinformatics analysis. CK, MG and KG performed experiments. RS, CB, LOC, ZN, GH, LCH and TBHG provided samples and commented on the manuscript. FT acquired funding and supervised studies in mice.

## METHODS

### Human tissue collection

Tissues were collected after obtaining informed consents from the subjects, with the approval of tissue-specific protocols by the Medical Ethical Committee of the Academic Medical Center (Amsterdam) and Erasmus MC (Rotterdam), following the Declaration of Helsinki protocols. Healthy skin tissues were obtained as excess materials from plastic surgery of the breast or abdomen. Intestinal samples were obtained after surgical resection with the exclusion of subjects that had undergone chemo- or radiotherapy prior to surgery. Fetal intestine was obtained from aborted fetuses at the Stichting Bloemenhoven clinic in Heemstede, the Netherlands. Gestational age ranged from 14 to 19 weeks, which was determined by ultrasonic measurement of the skull or femur diameter. Lung tissues were obtained from excess materials from lobectomy surgery of NSCLC at Erasmus MC Rotterdam. Nasal polyp samples were collected from patients with CRSwNP by endoscopic sinus surgery. Buffy coat was collected from healthy volunteers recruited by the blood bank at the Sanquin Blood bank. Asthma, COPD, Smoking and non-smoking control blood was obtained from the Sint Franciscus Gasthuis, Rotterdam.

### Isolation of cells from human samples

Single cell suspension was obtained from intestinal tissues (adult and fetal) and skin as previously described^22,36,37^. Nasal tissues were processed as preciously described^49^. Lung tissue was cut into small pieces, blood was washed away with PBS and tissue pieces were digested in RPMI + Collagenase D (1 mg/ml) + DNase I (0.1 mg/ml) for 1 hour. PBMCs were isolated from buffy coats as previously described^38^.

### Mice

C57BL/6J (B6) mice were bred in the British Columbia Cancer Research Centre (BCCRC) animal facilities from breeders obtained from the Jackson Laboratory. Rora-YFP mice were generated in-house as previously described^23^. All animal use was approved by the animal care committee of the University of British Columbia and mice were maintained in the BCCRC specific pathogen free facility. Briefly, mice were housed in vented cages (4 mice maximum per cage) with crinkle paper or cotton nesting materials and a hiding place. They were fed a normal chow diet. Mice were anesthetized by isoflurane inhalation during intranasal treatment to minimize animal suffering and distress. After the injections, they were moved to a separate cage placed on a heating pad and monitored closely until they were fully recovered from anesthesia. The health status of the mice was assessed daily during the period of treatments and one day after the last injections based on their appearance and behaviour. At the time of harvest, mice were anesthetized by isoflurane inhalation until they were unconscious and euthanized by carbon dioxide asphyxiation. All mice were used between 6–12 weeks of age.

### Tissue processing and primary leukocyte preparation from mice

Lungs were collected from mice and chopped into small pieces using a razor blade. Pieces of lungs were enzymatically digested in 5 mL of DMEM + 10% FBS, supplemented with 118.05 Kunitz U/mL Deoxyribonuclease I and 142.5 U/mL Collagenase IV (37 °C, 25 min, 250 rpm shaker). Digested lungs were mashed through 70 μm strainer and washed with 5 mL DMEM + 10% FBS, after which they were centrifuged (5 min, 400 xg) and the supernatant was discarded. Leukocytes were isolated by density gradient centrifugation (10 min, 650 xg, no acceleration or brake) in 5 mL of 36% percoll solution. Red blood cells were lysed in 3 mL of 150 mM ammonium chloride solution and washed with 5 mL PBS + 2% FBS.

For preparation of single cell suspension from the small intestine, Peyer’s patches were removed, and the intestine was fragmented into 3-4 pieces. Each piece was longitudinally cut open and washed in 5 mL wash buffer (88.5% water, 10% of 10x Hanks’ Balanced Salt solution and 1.5% of 1M HEPES, pH=7.2) until clean. The cleaned pieces were further cut into 1 cm and washed in the wash buffer by vigorously shaking three times, after which they were resuspended in 20 mL pre-warmed (37 °C) intestinal epithelial fraction buffer (77.4% water, 10 % of 10x HBSS, 10% FBS, 1.5% of 1 M HEPES, 1% of 0.5 M EDTA and 0.1% of 1 M dithiothreitol, pH=7.2) and incubated while shaking (37 °C, 20 min, 250 rpm shaker). The buffer was removed after vigorous shaking and samples were washed 3 times in wash buffer as above. The incubation step was repeated and the residual intestine pieces were resuspended in 20 mL lamina propria buffer (88.4% RPMI 1640 with HEPES and without glutamine, 10% FBS, 1.5% of 1 M HEPES, and 0.1% of 1 M DTT, pH 7.2), supplemented with 100 U/mL collagenase VIII (Sigma, C2139) and 9.41 Kunitz U/mL DNase I. Samples were incubated at 37 °C for 1 hour in a shaker at 250 rpm, after which the supernatant was collected through 70 μm strainer. The incubation step was repeated for the second time, and the cells were collected by centrifuging the combined flow through (4 °C, 15 min, 400 xg). Cells were resuspended in 5 mL 90% percoll, which was overlaid with 8 mL of 36% percoll, and subjected to density gradient separation (20 min, 650 xg, no acceleration or brake). Leukocytes were isolated by collecting the interphase and washed with 12 mL PBS + 2% FBS and pelleted by centrifugation (4 °C, 10 min, 400 xg).

For single cell suspension of skin cells, both ears were collected from mice and split into dorsal and ventral halves using forceps. Ear tissues were digested in 5 mL PBS + 2% FBS containing 420 U/mL type IV collagenase (37 °C, 40 mins, 250 rpm shaker). Digested skin was then mashed through 70 μm strainers, followed by washing with 5 mL PBS + 2% FBS. Cells were collected by centrifugation (4 °C, 5 min, 400 xg) and washed again with 10 mL PBS + 2% FBS.

### Antibodies, Reagents and Flow cytometers

Isolated leukocytes were counted using a hemocytometer and incubated in 2.4G2 mAb to block Fc receptors for analyses by fluorescence-activated cell sorting (FACS). Fluorescein isothiocyanate (FITC)-conjugated anti-mouse KLRG1 (2F1), PerCP-Cy5.5-conjugated anti-mouse CD25 (PC61.5), PerCP-eFluor 710-conjugated anti-mouse T1/ST2 (RMST2-2), rat IgG2a, κ isotype control (eBR2a), Allophycocyanin (APC)-conjugated anti-mouse KLRG1 (2F1), eFluor 660-conjugated anti-mouse/human GATA3 (TWAJ), AF700-conjugated anti-mouse CD127 (A7R34), eFluor 450-conjugated CD11b (M1/70), CD11c (N418), CD19 (1D3), CD3ε (145-2C11), CD4 (RM4-5), Gr-1 (RB6-8C5), NK1.1 (PK136), TCRβ (H57-597), TCRγδ (GL3), Ter119 (TER-119), PE-conjugated anti-mouse IL-13 (eBio13A), anti-mouse/human GATA3 (TWAJ), rat IgG2a, κ isotype control (eBR2a), PE-Texas Red-conjugated anti-human CD45RA (MEM-56), PECy7-conjugated anti-mouse CD127 (A7R34) were purchased from Thermo Fisher Scientific. APC-conjugated anti-mouse/human IL-5 (TRFK5), rat IgG1, κ isotype control (R3-34), BV421-conjugated anti-human CD200R (OX-108), V500-conjugated anti-mouse CD45 (30-F11), BV650-conjugated anti-human ICOS (DX29), PE-conjugated anti-human CD25 (M-A251) were purchased from BD Biosciences. FITC-conjugated anti-human CRTH2 (BM16), CD1a (HI149), CD3ε (OKT3), CD4 (RPA-T4), CD14 (HCD14), CD19 (HIB19), CD34 (581), CD123 (6H6), TCRαβ (IP26), TCRγδ (B1), FcεRIα (AER-37 (CRA-1)), BDCA2 (201A), CD56 (HCD56), AF700-conjugated anti-human CD3ε (UCHT1), APCCy7-conjugated anti-human CD45 (2D1), CD45RO (UCHL-1), Brilliant Violet (BV) 510-conjugated anti-human CD161 (HP-3G10), BV605-conjugated anti-mouse CD90.2 (53-2.1), PE-conjugated rat IgG1, κ isotype control (RTK2071), PE/Dazzle 594-conjugated anti-human CD200R1 (OX-108), PECy7-conjugated anti-human CD127 (A01905), rat IgG2a, κ isotype control (RTK2758) were purchased from BioLegend. FITC-conjugated anti-mouse anti-mouse T1/ST2 (DJ8) was purchased from MD Biosciences. BV605- conjugated anti-human CD45 (HI30) was purchased from Sony. PECy5.5-conjugated anti-human CD117 (104D2D1) was purchased from Beckman Coulter. BD LSRFortessa was used for flow cytometric analyses. Samples were first gated on live cells using eFluor 780 fixable viability dye, followed by single cell gating. Mouse lineage cocktail contains CD3ε, CD4, CD19, TCRβ, TCRγδ, CD11b, CD11c, NK1.1, Ter119 and Gr-1, while human lineage cocktail contains CD1a, CD3ε, CD4, CD5, CD14, CD16/32, CD19, CD34, CD123, TCRαβ, TCRγδ, FcεRIα, BDCA2, CD8α, CD11c, CD56. Flowjo version 10.0.7r2 was used for data analyses.

### In vitro culture

For in vitro culture of human ILC2s, purified cells were plated in 96-well round bottom plate in Yssel’s media (IMDM + 4% v/v Yssel’s supplement (made in house, Amsterdam UMC) + 1% v/v human AB serum (Invitrogen)). ILC2s were cultured in the presence of IL-2 (10 U/ml) plus IL-7, and different combinations of IL-25, IL-33, TSLP, IL-1β, IL-18 (all 20 ng/ml), specified in the figure legends. Cytokines were supplemented every 3 days. IL-5 was detected by intracellular staining and flow cytometry.

### Quantitative real-time PCR

Purified cells were lysed and RNA was isolated using the NucleoSpin RNA XS kit (Macherey-Nagel) according to the manufacturer’s protocol. Complementary DNA was synthesized using High-Capacity Archive kit (Applied Biosystems). qPCR was performed with LightCycler 480 Instrument II (Roche) with SYBR Green I master mix (Roche).

### CEL-Seq2-based single-cell RNA sequencing

For single-cell RNA sequencing, ILCs were purified from the dermis of healthy skin. Isolation was performed directly after surgery. 1583 cells were directly sorted in 384-well plates, frozen and sent to Single Cell Discoveries for further processing of their pipeline based on CEL-Seq2^19^. Libraries were run on a HiSeq4000 for Illumina sequencing. Post-processing and quality control were performed by Single Cell Discoveries. Reads were aligned to GRCh38 reference assembly using STARsolo^39^. All transcriptome data analysis was performed using Seurat package v4. Adjusted P values were calculated using Bonferroni correction.

### Single cell cloning

3000 per well of OP9-DL1 murine stromal cells were plated in 96-well round-bottom plate at least 5 hours prior to the start of the culture. CD127^+^ and CD127^-^ ILC2s were sorted as single cells per well in the 96-well round-bottom plate and were expanded in Yssel’s medium supplemented with 1% (vol/vol) human AB serum in the presence of IL-23, IL-1β and TGF-β (all at a concentration of 20ng/ml). Clones were analyzed within 14 days of culture.

### Cell staining for FACS analyses

Cells isolated from tissues were stained with surface stains for half an hour at 4 °C, followed by FACS analyses or intracellular staining. For intracellular cytokine staining of murine samples, leukocytes were incubated at 37°C for 3 hours in 500 µL RPMI 1640 media containing 10% FBS, 100 U/mL P/S, 50 μM 2-ME, Brefeldin A (Golgi Plug, BD Biosciences), 30 ng/mL phorbol 12-myristate 13-acetate (PMA, Sigma, P1585) and 500 ng/mL ionomycin (Sigma, 10634). Human cells were stimulated 3 hours with PMA (10 ng/mL) and ionomycin (500 nM), with Golgi Plug in the last 5 hours of the culture. Intracellular cytokine staining was performed after the pre-incubation and surface staining, using Cytofix/Cytoperm Fixation/Permeabilization Solution kit (BD Biosciences) according to manufacturer’s protocol. Transcription factor staining was performed without pre-incubation using Foxp3/Transcription Factor Staining Buffer Set (Thermo Fisher Scientific) according to manufacturer’s protocol. For intracellular cytokine and intranuclear staining of Rorα-YFP mice, cells were pre-fixed with 0.5 % paraformaldehyde for 5 minutes at room temperature, before proceeding to fixation, permeabilization and intracellular staining.

### In vivo stimulation of mice

Mice were anesthetized by isoflurane inhalation and given i.n. administrations of 0.25 µg IL-33 (BioLegend) or 0.218U papain (Sigma) in 40 µL PBS.

## SUPPLEMENTARY INFORMATION TITLES AND LEGENDS

**Supplementary figure 1.**
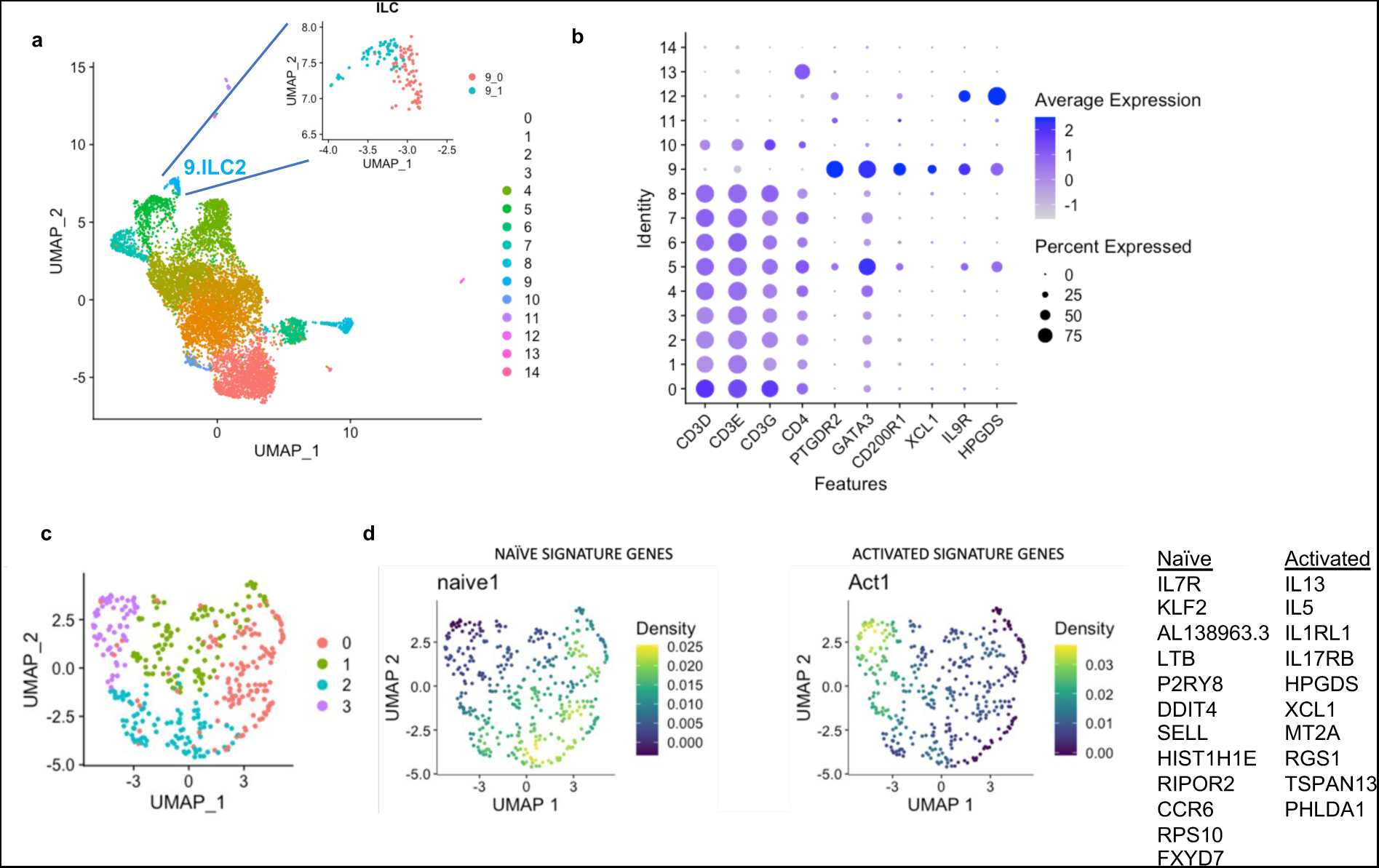
**a)** UMAP visual representation of the T and ILC2 (Lin- CRTH2+) clusters obtained from the analysis of the scRNAseq dataset from nasal polyps from CRSwNP patients^17^. **b)** Dot plot showing featured genes for each cluster identified in a). **c)** UMAP visual representation of the cell clusters obtained from the analysis of the scRNAseq dataset from atopic dermatitis biopsies^18^. **d)** Density plots of signature genes of naïve and activated ILC2s from the scRNAseq dataset from atopic dermatitis biopsies.

**Supplementary figure 2.**
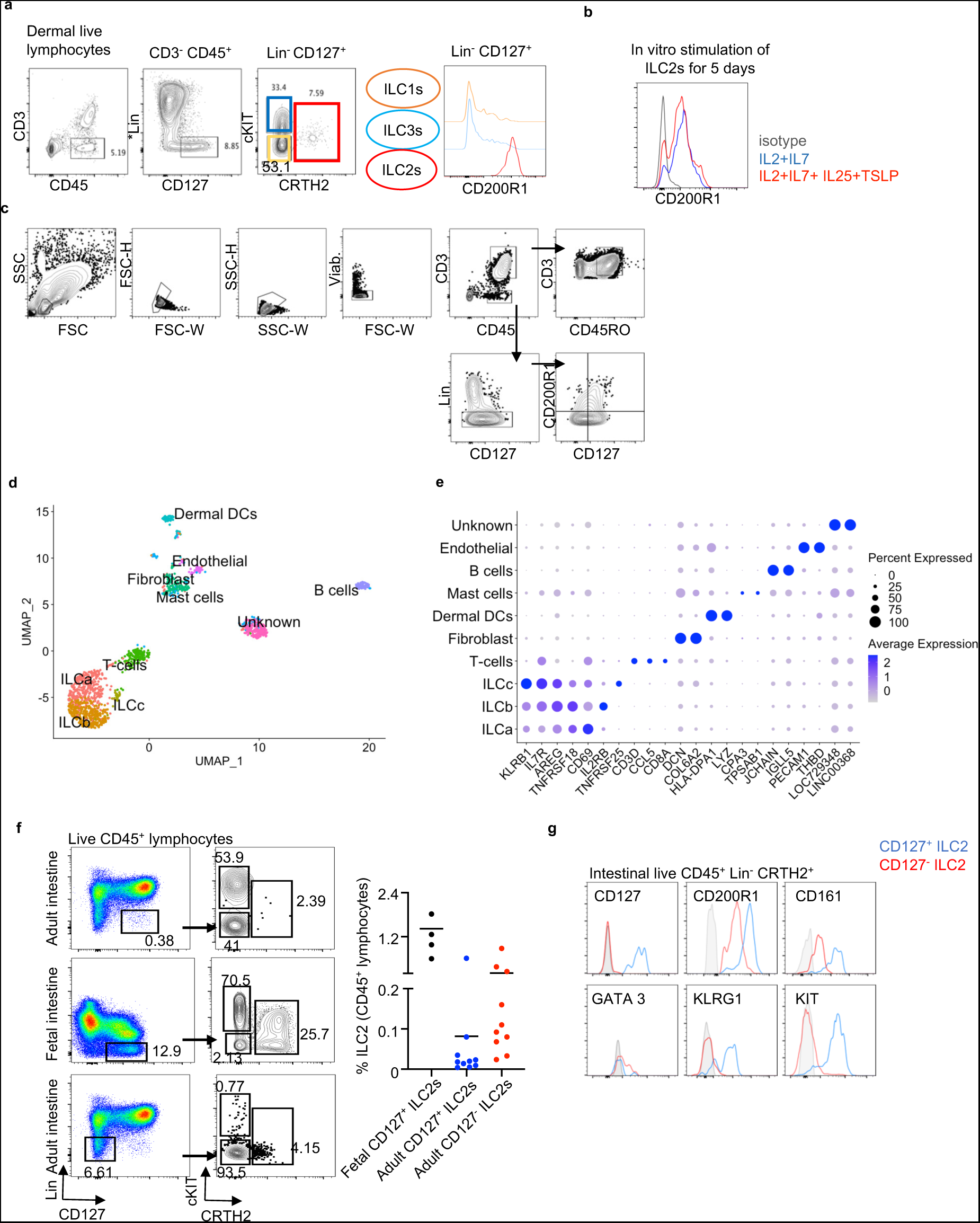
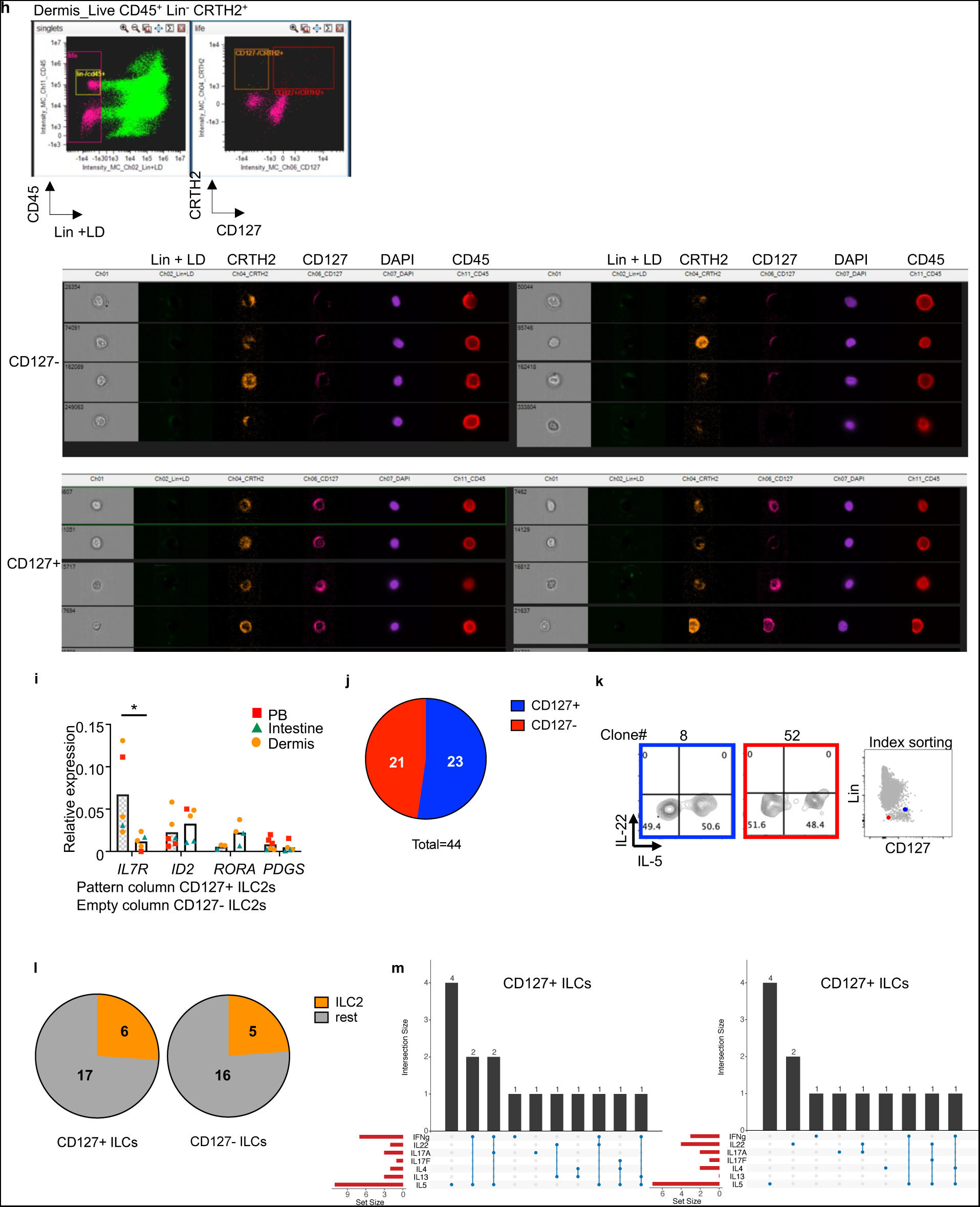
**a)** Full gating strategy and expression of CD200R1 of healthy human dermal ILCs analyzed by flow cytometry. **b)** Flow cytometry measurement of CD200R expression of in vitro stimulated PB ILC2s for 7 days with IL-2+IL-7+IL-25+TSLP or IL-2+IL-7 as control for 7 days. **c)** Gating strategy for sorting of human dermal Lin- cells including ILCs for the single cell RNA sequencing experiment. Cells were gated on viable lymphocytes CD45+ Lin- (CD1a, CD3, CD5, CD8, CD11c, CD14, CD16, CD19, CD34, CD94, CD123, FceR1a, TCRαβ, TCRgd, BDCA2) and then gated in 4 quadrants using CD127 and CD200R1 expression. Equal numbers of the 4 populations were sorted. **d)** UMAP visual representation of the cell clusters obtained from the scRNAseq analysis of Lin- cells sorted from healthy dermis. Clusters were calculated using Seurat. **e)** Dot plot representing the expression of marker genes from each cluster identified in the scRNAseq analysis. The intensity of the color indicates the level of expression of the indicated gene. The size of the symbol indicates the percentage of cells within the cluster expressing the corresponding marker gene. **f)** Flow cytometry analysis of CD127+ and CD127- ILC2s in adult and fetal intestine. **g)** Flow cytometry measurement of the expression of CD127, CD200R, CD161, GATA3, KLRG1 and cKIT on CD127+ and CD127- ILC2s from adult intestine. **h)** ImageStream Flow cytometry of CD127+ and CD127- ILC2s from dermal cells. **i)** Quantitative PCR of the expression of *IL7R*, *ID2*, *RORA* and *PDGS* of CD127+ and CD127- ILC2 isolated from PB, intestine and dermis. **j)** Pie chart showing the number of total clones from CD127+ and CD127- ILCs obtained after single cell sorting of ILCs from PB. **k)** Flow cytometry measurement of IL-5 expression from ILC2 clones (left) and index sorting phenotype of the clones (right). **l)** Pie chart showing the number of ILC2 clones from CD127+ and CD127- ILCs. **m)** Summary of numbers and type of cytokines produced by CD127+ and CD127- ILC clones. Data from a), b) and g) are representative of at least three donors from more than three independent experiments. Symbols represent individual donors; bars indicate mean values. *P < 0.05, (Paired t test).

**Supplementary figure 3.**
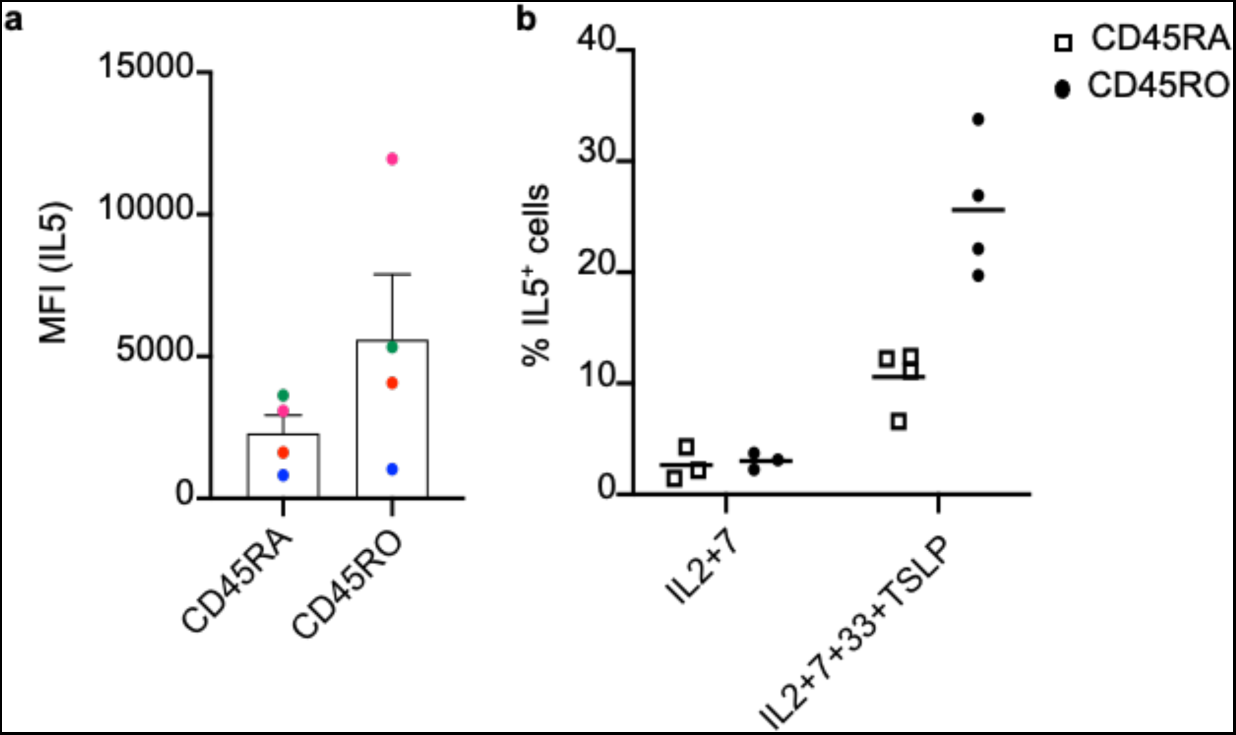
**a)** Mean fluorescence intensity (MFI) for IL-5 of CD45RA+ and CD45RO+ ILC2s isolated from PB and in vitro stimulated with IL-2 + IL-7 + IL-33 + TSLP for 24h. **b)** Percentage of IL-5 expression on CD45RA+ and CD45RO+ dermal ILC2s after 24h stimulation with IL2+IL7+IL33+TSLP or IL2+IL7 as control.

**Supplementary figure 4.**
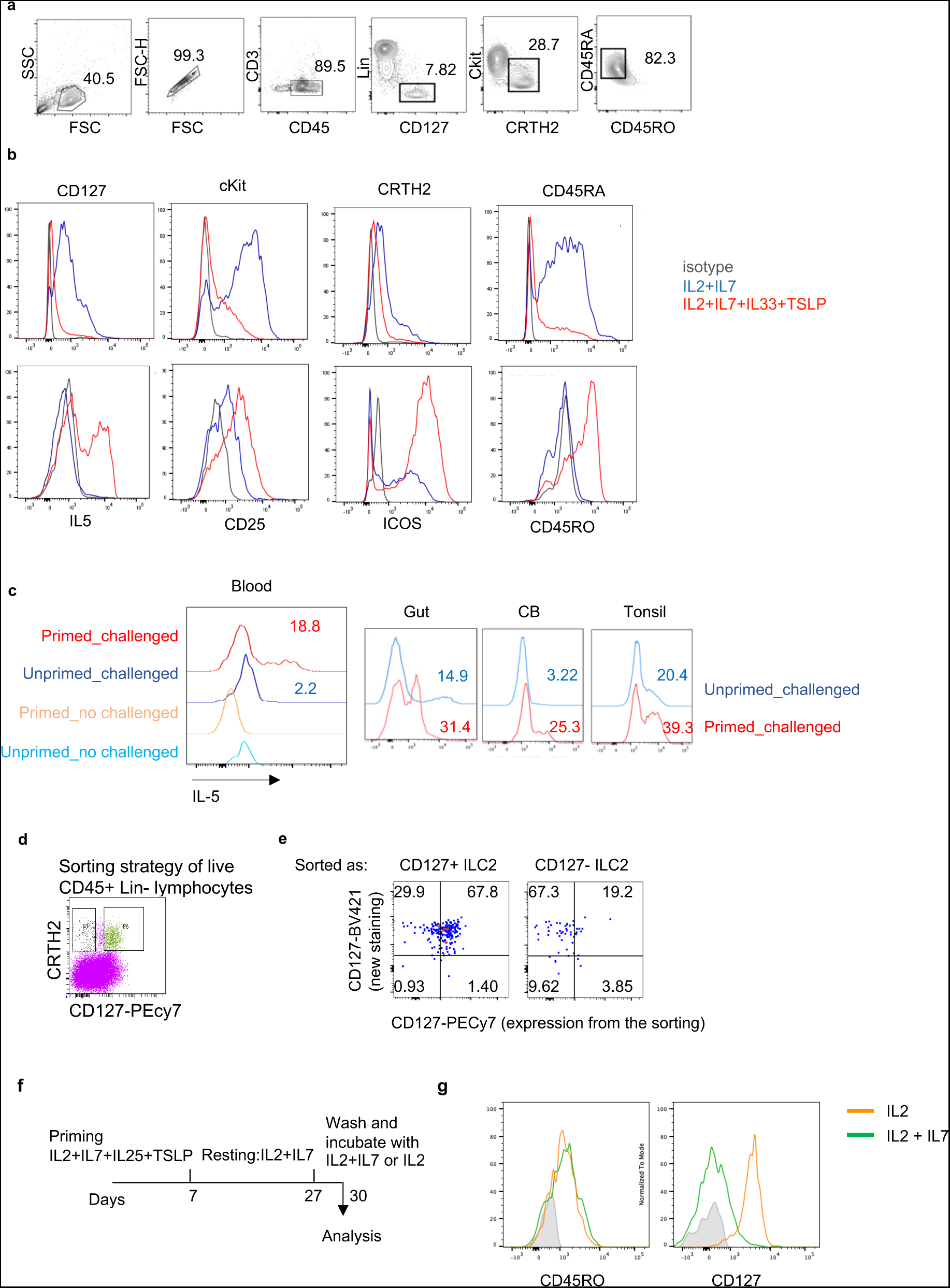
**a)** Gating strategy for ILC2 sorting from PB (as an example) for the long-term in vitro culture experiment. **b)** Flow cytometry measurements of the expression of the indicated markers on in vitro stimulated (red) and control (blue) ILC2s on day 7. **c)** Flow cytometry measurement of IL5 expression of challenged (primed or unprimed) ILC2s isolated from the indicated tissues. **d)** Sorting strategy of CD127+ and CD127- ILC2s from healthy dermis for the short-term in vitro culture in the presence of IL-2. **e)** Flow cytometry analysis of CD127 expression on CD127+ and CD127- isolated dermal ILC2s and cultured in the presence of IL-2 for 24h. CD127 was stained with an antibody labelled with a different fluorochrome but same clone than the one used for the sorting. **f)** Scheme of the long-term in vitro culture with IL7 withdrawal. **g)** Histogram of the expression of CD45RO and CD127 on the cultured ILC2s in the indicated conditions. Data are representative of at least three donors from more than three independent experiments. Symbols represent individual donors; bars indicate mean values.

